# Computing wide range of protein/peptide features from their sequence and structure

**DOI:** 10.1101/599126

**Authors:** Akshara Pande, Sumeet Patiyal, Anjali Lathwal, Chakit Arora, Dilraj Kaur, Anjali Dhall, Gaurav Mishra, Harpreet Kaur, Neelam Sharma, Shipra Jain, Salman Sadullah Usmani, Piyush Agrawal, Rajesh Kumar, Vinod Kumar, Gajendra P.S. Raghava

## Abstract

**Motivation:** In last three decades, a wide range of protein descriptors/features have been discovered to annotate a protein with high precision. A wide range of features have been integrated in numerous software packages (e.g., PROFEAT, PyBioMed, iFeature, protr, Rcpi, propy) to predict function of a protein. These features are not suitable to predict function of a protein at residue level such as prediction of ligand binding residues, DNA interacting residues, post translational modification etc.

**Results:** In order to facilitate scientific community, we have developed a software package that computes more than 50,000 features, important for predicting function of a protein and its residues. It has five major modules for computing; composition-based features, binary profiles, evolutionary information, structure-based features and patterns. The composition-based module allows user to compute; i) simple compositions like amino acid, dipeptide, tripeptide; ii) Properties based compositions; iii) Repeats and distribution of amino acids; iv) Shannon entropy to measure the low complexity regions; iv) Miscellaneous compositions like pseudo amino acid, autocorrelation, conjoint triad, quasi-sequence order. Binary profile of amino acid sequences provides complete information including order of residues or type of residues; specifically, suitable to predict function of a protein at residue level. Pfeature allows one to compute evolutionary information-based features in form of PSSM profile generated using PSIBLAST. Structure based module allows computing structure-based features, specifically suitable to annotate chemically modified peptides/proteins. Pfeature also allows generating overlapping patterns and feature from whole protein or its parts (e.g., N-terminal, C-terminal). In summary, Pfeature comprises of almost all features used till now, for predicting function of a protein/peptide including its residues.

**Availability:** It is available in form of a web server, named as Pfeature (https://webs.iiitd.edu.in/raghava/pfeature/), as well as python library and standalone package (https://github.com/raghavagps/Pfeature) suitable for Windows, Ubuntu, Fedora, MacOS and Centos based operating system.

## 1. Introduction

Advent of next generation sequence technologies earmarked the production of a huge amount of genomic and proteomic data that leads to exponential growth of protein databases. The experimental techniques used for functional annotation of a protein are time consuming, costly and slow. Thus, functional and structural annotation of a protein is one of the main tasks in the field of bioinformatics. In past, several hundred methods have been developed for predicting different type of protein properties. Broadly, these methods can be divided in three categories; i) function at protein level, ii) residue level annotation and iii) therapeutic annotation of chemically modified peptides. In all these methods, one of the major challenges is to depict numerical representation a protein, which is known as protein feature or descriptor, although last five decades have witnessed significant progress in detecting new protein features. Functional annotation of a protein can be achieved by predicting overall function of a protein which include subcellular localization, classification and diversified properties of a protein. Subcellular localization of a protein is vital in understanding its function, TargetP (Emanuelsson *et al.*, 2000) is one of the initial methods developed for subcellular localization beside other significant methods such as ESLpred (M. Bhasin and Raghava, 2004a), WoLF (Horton *et al.*, 2007), PSLpred (Bhasin *et al.*, 2005), HSLpred (Garg *et al.*, 2005), RSLpred (Kaundal and Raghava, 2009), MultiLoc2 (Blum *et al.*, 2009). Similarly method have been developed to specific class of proteins that includes GPCRpred (M. Bhasin and Raghava, 2004b), NRpred (Manoj Bhasin and Raghava, 2004), CytoPred (Lata and Raghava, 2008), CyclinPred (Kalita *et al.*, 2008).

These methods utilize composition-based features to represent a variable length of a protein by fixed length vector. For example, amino acid composition-based feature represents a protein of variable length by a vector of dimension twenty. Numerous composition-based features such as pseudo amino acid, autocorrelation, conjoint triad, quasi-sequence order etc. have been reported over the year. Likewise, several methods to predict therapeutic properties of a protein or peptide such as anti-cancer, anti-microbial, anti-bacterial, anti-fungal, anti-tubercular, anti-hypertensive, toxic, tumor homing etc. have been developed (Manavalan *et al.*, 2017; Tyagi *et al.*, 2013; Meher *et al.*, 2017; Lata, Mishra, and G. P. S. Raghava, 2010; Agrawal *et al.*, 2018; Usmani, Bhalla, *et al.*, 2018; Kumar *et al.*, 2015; Manavalan *et al.*, 2018; Gupta *et al.*, 2013; Sharma *et al.*, 2013). These peptides/proteins have therapeutic properties thus play vital role in designing proteins/peptide-based drugs and vaccines (Dhanda *et al.*, 2017; Usmani, Kumar, *et al.*, 2018; Nagpal *et al.*, 2017). Last few decades witnessed significant increase in the FDA approval of peptide-based drugs (Usmani *et al.*, 2017). In addition to composition of a whole peptide, composition of N-terminal or C-terminal residues of peptides have also been used for prediction. Most of the natural peptides have limited half-life in body fluids and in intestine. These peptides are chemically modified to improve their stability, most of FDA approved drugs are chemically modified. Thus, computing descriptors of these chemically modified peptides is a challenge as amino acid sequence information is not adequate to represent these peptides. In past methods like AntiMPmod (Agrawal and Raghava, 2018), CellPPDmod (Kumar *et al.*, 2018) have been developed for predicting therapeutic properties of chemically modified peptides. In these methods, structural and chemical based descriptors have been used for developing prediction models.

Although, protein annotation at sequence level provides overall property of a protein but residue level annotation is important to understand function of a protein (Nagel *et al.*, 2009). One of the initial methods developed for predicting secondary structure state of each residue in a protein was Chou-Fasman method. It utilizes the combination of statistical and heuristic approach in order to predict the secondary structure state of protein residues. It’s successful implementation encourages researchers to use residue level features in predicting irregular secondary structure like alpha turns (Wang *et al.*, 2006), beta turns (Singh *et al.*, 2015; H. Kaur and Raghava, 2003; Fuchs and Alix, 2005; Kountouris and Hirst, 2010), gamma turns (Guruprasad and Rajkumar, 2000; Jahandideh *et al.*, 2007), beta hairpins (de la Cruz *et al.*, 2002) and beta barrel (Freeman and Wimley, 2012). Initially, these methods were developed using binary profile of protein patterns/segments where a residue is represented by a vector of 20 dimensions. A pattern of length 17 residues will be represented by a vector of dimension 340 (17 × 20). The performance of these methods improved significantly when evolutionary information has been used instead of protein sequence. In order to utilize evolutionary information, protein profiles were generated to capture evolutionary information from similar sequences. One of the commonly used protein profile is PSSM profile which is generated using PSI-BLAST (McGinnis and Madden, 2004). PSSM profiles have been heavily used as feature for predicting function of a protein at residue level.

In past, numerous web servers, standalone software and libraries has been developed that includes PROFEAT (Li *et al.*, 2006), PyBioMed (Dong *et al.*, 2018), iFeature (Chen *et al.*, 2018), protr (Xiao *et al.*, 2015), Rcpi (Cao *et al.*, 2015), propy (Cao, Xu, *et al.*, 2013); for computing features of a protein. Despite number of methods have been developed in past, still number of important features have not been integrated in any software, particularly features required for annotating protein at residue level. In order to complement previous efforts, we make a systematic attempt to provide platform which integrate most of features discovered in past.

## 2. Materials and Methods

In this study, we have mainly implemented existing algorithms for computing different type of features. In addition to existing algorithms, we have also developed algorithms for specific type of features. Though these features have not been tested in model’s but provide important information about a protein. In the following section, summary of these algorithms has been briefly described.

### 2.1. Composition-based Features

These features provide overalls content of a protein like amino acid composition where fraction of each type of residue is computed from amino acid sequence. This type of numerical representation of a protein is important for developing models using machine learning techniques. One of the major advantage is their ability to represent variable length protein by a fixed number of features. Following is brief description of these features, while detail description is given in manual (https://webs.iiitd.edu.in/raghava/pfeature/Pfeature_Manual.pdf).

#### 2.1.1. Simple Compositions

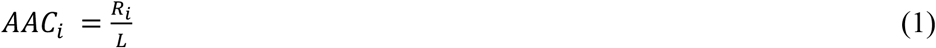

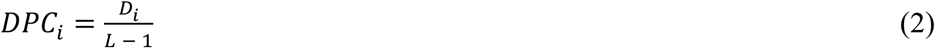

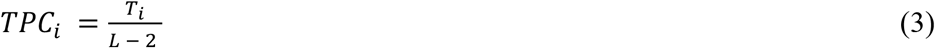

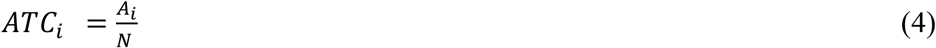

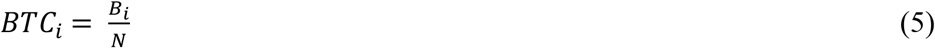

where *AAC*_*i*_, *DPC*_*i*_, *TPC*_*i*_, *ATC*_*i*_ and *BTC*_*i*_ are composition of residue, dipeptide, tripeptide, atom and bond composition of type *i* respectively. *R*_*i*_, *D*_*i*_, *T*_*i*_, *A*_*i*_ and *B*_*i*_ are number of residues, dipeptides, tripeptides, atoms and bonds of type *i* respectively. *L* is length of protein, *N* is total number of atoms in case of atom composition and number of bonds in case of bond composition.

#### 2.1.2. Physico-Chemical properties

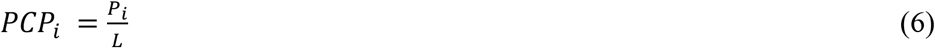

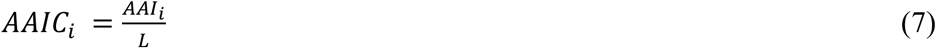

where *PCP*_*i*_ and *AAIC*_*i*_ are composition of physico-chemical properties and AAindex of residue type *i* respectively. *P*_*i*_ and *AAI*_*i*_ are sum of physico-chemical properties and AAindex values of residue type *i* respectively. L is length of protein sequence.

#### 2.1.3. Repeats & Distribution

Simple composition and physico-chemical property based modules measure fraction of residue and their property. One of the problem with existing features is that they do not measure repeat of particular type of residue or distribution. In this study, we introduced new features, which compute repeats of amino acids and distribution of amino acids.

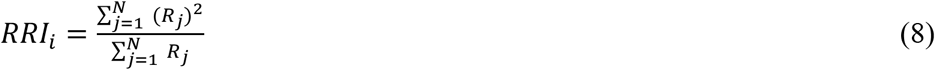

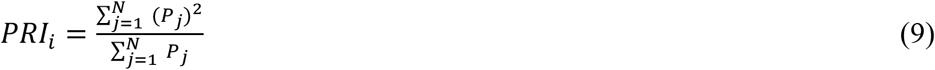

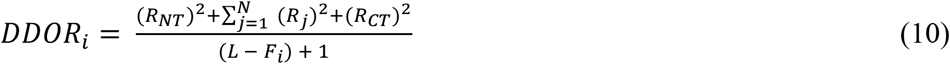

where *RRI*_*i*_, *PRI*_*i*_ and *DDOR*_*i*_ are residue repeat, property repeat information and distance distribution of residue type ***i*** respectively. *N, R*_*j*_ and *P*_*j*_ are maximum number of occurrences, number of runs/repeats of residue type *j* and number of runs/repeats of property type *j* respectively. *R*_*NT*_, *R*_*j*_, *R*_*CT*_, *L* and *F*_*i*_ are residue distance from N-terminal, inter-distance between residue type *i*, residue distance from C-terminal, total length of protein sequence and frequency of residue type *i* respectively.

#### 2.1.4. Shannon Entropy

In order to measure level of complexity at protein and at residue level, we compute Shannon entropy of a protein and entropy of each type of residues using following equations

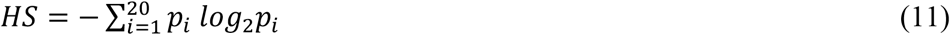

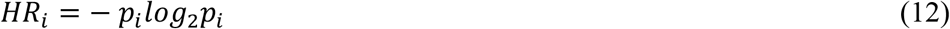

where *HS* is Shannon entropy of a protein sequence and *HR*_*i*_ is entropy of a residue type *i*. *p*_i_ is the probability of a given amino acid in the sequence. This equation was extended to compute entropy of a particular type of property like charge, polarity, hydrophobicity in a protein sequence.

#### 2.1.5. Miscellaneous compositions

In past wide range composition-based features have been generated from proteins. We implement these features in Pfeature using algorithm described in previous studies (**References**). These features includes; Autocorrelation, Conjoint Triad Descriptors, Composition enhanced Transition and Distribution, Pseudo Amino Acid Composition, Amphiphilic Pseudo Amino Acid Composition, Quasi-Sequence Order, Sequence Order Coupling Number.

#### 2.1.6. PSSM Composition

In past evolutionary information as PSSM composition have been used for classification of proteins particularly for subcellular localization of proteins (Rashid *et al.*, 2007; Kumar *et al.*, 2007, 2011). In PSSM profile, the evolutionary information is presented by a matrix with dimension L × 21 (L rows and 21 columns) for a L length protein. We have generated vector of dimension 420 called PSSM-420, which maintain composition (Kumar *et al.*, 2007).

### 2.2. Binary Profiles

One of the major limitations of composition-based features is that they only maintain content of protein but not order of amino acids. Thus, composition-based features alone, are not suitable for predicting function of protein residues. In order to predict or annotate function of a protein at residue level, we need to compute features that also maintain order of amino acids in sequences.

#### 2.2.1. Amino Acid Profile

In past, binary profiles have been widely used for residue level annotation that includes prediction of protein’s secondary structure as well as nucleotide or ligand binding sites in a protein (H. Kaur and Raghava, 2003; Harpreet Kaur and Raghava, 2003, 2004; H. Kaur and Raghava, 2004; Kumar *et al.*, 2007; Ansari and Raghava, 2010; Panwar *et al.*, 2013). We have integrated binary profile in Pfeature and the details about binary profile is already described in previous studies (Singh *et al.*, 2015; Chauhan *et al.*, 2013, 2012), following is brief procedure for creating binary profile. First, overlapping fixed length patterns are generated from protein sequences. Secondly, each pattern (or peptide segment) will be presented numerical numbers. Each amino acid will be presented by 21 numerical values, 20 units for 20 amino acids and one unit for adding dummy amino acid at both terminus to create one pattern corresponding to each number. Each amino acid is presented by a vector of dimension 21, if amino acid is present then value of element is 0 otherwise one. For example A is presented by vector (1,0,0,0,0,0,0,0,0,0,0,0,0,0,0,0,0,0,0,0,0), C by (0,1,0,0,0,0,0,0,0,0,0,0,0,0,0,0,0,0,0,0,0), D by (0,0,1,0,0,0,0,0,0,0,0,0,0,0,0,0,0,0,0,0,0) and so on. Thus a pattern of length will be presented by vector of dimension N × 21.

#### 2.2.2. Dipeptides Profile

In this study, we have introduced new features called dipeptide profiles, similar to amino acid profiles. We have created a vector of dimension 400 for 20 amino acids and each dipeptide is represented by a vector of dimension 400, where presence of dipeptide is represented by one and absence by 0. This feature has not been tested so far in any prediction method but we belive, it can be used in future studies.

#### 2.2.3. Atom and Bond Profile

Recently, a method has been developed for predicting antimicrobial activity of chemically modified peptides (Agrawal and Raghava, 2018). In this study, atom and bond profiles have been used for creating features for chemically modified peptides. This atom and bond-based features, provide more precise information than amino acid. In the case of atoms profile, there were total 5 atoms (C, H, O, N, S) where the presence of atom was represented by “1” and the absence by “0”, hence generating a vector of N × 5. Their combinations form binary profile of each residue. For example, residue of Alanine (A) contains 13 atoms in total. Thus, binary profile of ‘A’ will be of size 13*5=65. Similarly, bonds profiles have been computed from protein sequences.

#### 2.2.4. Residue Properties Profile

Here, we have represented a protein by physico-chemical properties of its residues. For example, if user need to create property-profile for positive charge residues. We will present an amino acid by “1” if it is positive charge otherwise “0”. This way, we may create properties profile for 25 type of physico-chemical properties like hydrophobicity, hydrophilicity, polarity. For example, hydrophobicity profile for amino acid sequence “DPARAAGAHQ” will be {0,1,1,0,1,1,0,1,0,0) as amino acid “P” and “A” are hydrophobic.

#### 2.2.5. AAindex Profile

Here, we have used AAindex values for computing binary profiles. If users, select 10 AAindex for creating profile then we present an amino acid with 10 values, each value will represent a one AAindex value of an amino acid. This function gives the binary profile of input AA Indices. If normalized score of AAIndex value of a particular residue is negative, the function assigns ‘0’ to that residue otherwise assigns ‘1’. For example binary profile for amino acid “A” for following AAindex values (ARGP820101, ANDN920101, ARGP820102, ARGP820103, BEGF750101, BEGF750102, BEGF750103, BHAR880101, BIGC670101, BIOV880101) will be {0,0,1,1,1,1,0,0,0,0}.

### 2.3. Evolutionary Information in form of PSSM Profile

Prediction based on evolutionary information are more accurate in comparison to the methods based on single profile. Here, evolutionary information is extracted in form of PSSM profiles, generated by PSIBLAST. Pfeature allows one to generate PSSM based features called PSSM profile. In past number of techniques have used to normalize PSSM, here we used following techniques for normalization (Kumar *et al.*, 2007).

- **pssm_n1**: Due to the large number of variation in the value of PSSM matrix, it is necessary to normalize it. Each element of matrix is normalized by 1/(1+e-x).
- **pssm_n2**: This is the second technique to normalize the elements of PSSM matrix using the formula (num - min)/(max - min).
- **pssm_n3**: This is the third technique to normalize the matrix using the formula (num - min)*100/(max - min).
- **pssm_n4**: This is the fourth technique to normalize the PSSM profile using the formula 1/(1+e-(x/100).

The PSSM contained the probability of occurrence of each type of amino acid residues at each position along with insertion/deletion. It measures conservation of a residue in a given location in protein. This means that evolutionary information for each amino acid is encapsulated in a vector of 21 dimensions where the size of PSSM matrix of a protein with N residues is 21×N. Thus, binary profile of a pattern/peptide of 9 amino acids will be dimension of 189 (21×9).

### 2.4. Features from portions of a sequence

In most of the protein classification methods, features are computed from whole protein. In literature, it has been shown that splitted amino acid composition (SAAP) based method perform better than composition-based methods (Restrepo-Montoya *et al.*, 2009; Kumar and Raghava, 2009; Garg and Raghava, 2008). In SAAP, proteins are splitted in number of parts and then the composition of each part is computed. This allows computing feature of different parts of proteins such as terminal, split and rest.

#### 2.4.1. N-Terminal

It has been observed in past that N-terminal of a protein is responsible for its function. For example, most of classical secretory proteins contain a signal peptide. A short peptide (16-30 amino acids) present at the N-terminus of the majority of proteins that are destined towards the secretory pathway. Pfeature allows user, to compute wide range of features in selected region (N-terminal) of a protein. User can generate both composition as well as binary profile as length of selected region is fixed (Lata *et al.*, 2007; Lata, Mishra, and G. P. Raghava, 2010).

#### 2.4.2. C-Terminal

The most common endoplasmic reticulum retention signal is the amino acid sequence KDEL or HDEL at the C-terminus. This keeps the protein in the endoplasmic reticulum and prevents it from entering the secretory pathway. Pfeature allow user to compute wide range of features in selected region (C-terminal) of a protein. One of the advantage of in selecting region is that user can generate both composition as well as binary profile as length of selected region is fixed (Lata, Mishra, and G. P. Raghava, 2010; Lata *et al.*, 2007).

#### 2.4.3. Split

In order to increase number of features to capture more information from a protein, split amino acid composition (SAAC) has been introduced (Kumar *et al.*, 2006). In this concept, amino acid is splitted in two or more than two portions then feature of each portion is computed separately. For example, if number of splits is three then sequence will be divided in three portions (each portion have nearly same length). If whole protein has 20 (composition) features then splitted composition provides 60 (20 × 3) features.

#### 2.4.4. Rest

As shown in above section both terminals (N- & C-) have important information so Pfeature have provision to compute feature of N-terminal or C-terminal. In order to capture information or generating feature from remaining portion of proteins (after removing N-terminal and C-terminal residues), rest option is provided, where user need to select number of residues from N-terminal and C-terminal to be removed.

## 3. Results

Aim of Pfeature is to facilitate users in computing most of existing features discovered in literature. It can be used as web server, where user can submit their sequence or structure on our web site for computing features. In addition, python scripts and libraries have been developed, so user can compute feature on their local machine. Following is brief description of different type of implementation of Pfeature and facilities.

### 3.1. Web Implementation

A web server Pfeature has been developed on Ubuntu operating system using Apache software. Most of web pages has been developed using HTML, PHP5 and CSS3. Submission page allow users to submit protein sequences in FASTA format or amino acid sequence of a protein in single line. In order to facilitate users, we provide example protein sequences. In addition to display results as HTML page, server allow to download results in csv format. Pfeature contain five major modules/parts, following is brief description (Figure 1).

**Figure 1:**
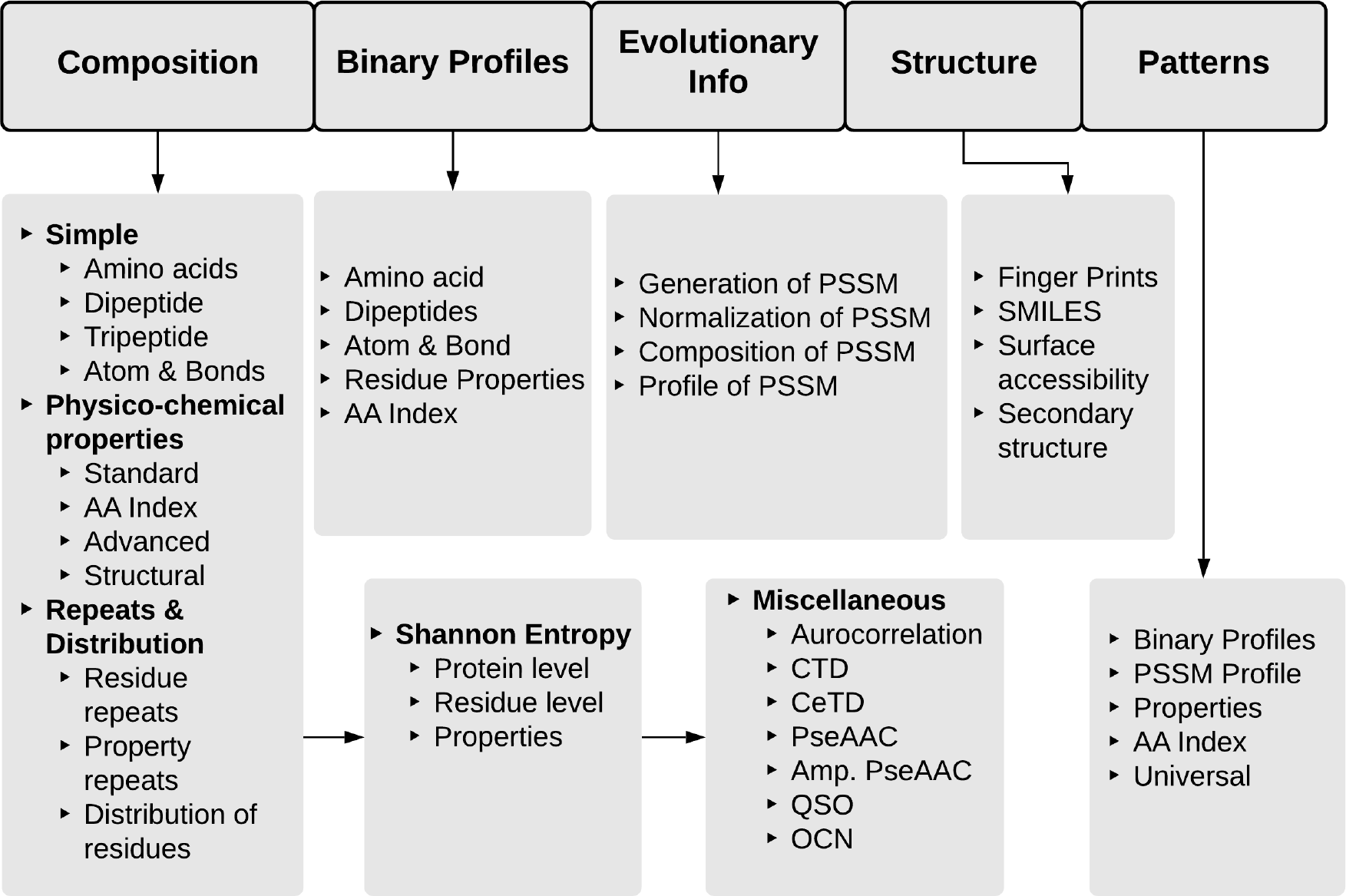
Overview of major features that can be calculated using Pfeature.

#### 3.1.1. Composition Module

This module is designed for computing wide range of composition-based features. These features have been classified in following five submenus/modules.

##### Simple Composition

This module allows users to compute simple composition like amino acid composition, dipeptide composition and tripeptide composition. In addition to compute different type of composition of whole sequence, Pfeature also allow to compute composition of a selected regions of a protein. This server also allows one to compute atom, bond and both type of composition in a protein. This is an interesting feature which allow to capture chemical modifications in a protein which is not practically possible from amino acid composition. It takes amino acid sequences in single-line format and gives output of some vector size depending upon the selection of the sub-module. In case of dipeptide composition, server also allows to compute higher order of dipeptide composition, which was introduced by our group (Garg *et al.*, 2005).

##### Physico-Chemical properties

This is similar to amino acid composition, here we compute composition of group of residues that have particular physico-chemical properties. It has following four sub menus; i) standard properties, ii) AAindex, iii) advanced and iv) structural. In case of standard physico-chemical properties, server compute composition of around 20 type of groups, each group having unique type of residues like, positive charged, polar, basic, acid residues. In case of AAindex, Pfeature allows user to compute composition of any amino acid index out of total 566 in database AAindex version 9.0. Advanced sub menu, has been developed for computing advanced physico-chemical properties like z1, z2, z3. This module allows to compute composition of advanced properties like secondary structure and surface accessibility of a protein sequence.

##### Repeats & Distribution

It is interesting menu or module of server that allows, measurement of repeats and distribution of residues in a protein. It has following major sub menus, i) RRI which in-turn is divided into five sub-menus as RRI (Residue Repeat Information), RRI_NT, RRI_CT, RRI_RT and RRI_ST ii) Property Repeats (PRI) iii) DDOR (Distance Distribution of Repeats).. It takes amino acid sequences in single-line format and gives output of some vector size depending upon the selection of the sub-module.

##### Shannon Entropy

This module is used to compute the Shannon information of protein/peptide sequences. It has three major parts, i) Shannon entropy at protein level ii) Shannon entropy at residue level iii) Shannon entropy of physico-chemical properties. It takes amino acid sequences in single-line format and gives output of some vector size depending upon the selection of the sub-module.

##### Miscellaneous

This part basically covers already existing methods of protein/peptide feature extraction (Dong *et al.*, 2018). This module have seven sub modules to compute complex composition based features, following is brief description of each features. Autocorrelation compute distribution of amino acid properties in the protein/peptide sequences. In case of Conjoint Triad Descriptors (CTD), amino acids are divided in seven groups then properties of triad are computed, output vector have dimension 343 (7×7×7). Composition enhanced Transition and Distribution (CeTD) compute enhance transition of residues after grouping. Pseudo Amino Acid Composition (PAAC) is one of the most widely used protein features heavily used by scientific community in past by Chou group (Chou, 2001). Amphiphilic Pseudo Amino Acid Composition (APAAC) is improved version of PAAC (Chou, 2005). Quasi-Sequence Order (QSO) descriptors are computed from the distance matrix between the 20 amino acids. Sequence Order Coupling Number (SOCN) allow user to compute a set of sequence-order-coupling numbers (Chou, 2000).

#### 3.1.2. Binary Profiles

This module is mostly used in annotating protein at residue level. A vector consisting of 0 and 1 can be generated depending upon the selection of sub-modules. The presence of particular residue is denoted by 1 and its absence is denoted by 0. The module is further subdivided into five sub-sections.

##### Amino acid

This sub-module permits users to generate the binary profiles corresponding to each residue of the peptide or protein sequences. This module also facilitates the generation of binary profiles of the subsets of the sequences such as from N-terminal, C-terminal, rest and split.

##### Dipeptides

This sub-module calculates the dipeptide binary profile with option to select the order of dipeptide. In case a protein has *N* residues, then number of traditional overlapping dipeptides will be *N-1*, it can be presented by a vector of dimension *(N-1)× 400*.

##### Atom & Bond

This module is sub-divided into three sections as Atom, Bond and Atom & Bond, which generates the binary profiles for atoms, bonds and both respectively. In case of atoms a vector will be represented by 5 based on five types of atom (C, H, N, O, S). In case of bonds, vector dimension is 4 based on four type of bonds used in this study (cyclic, benzene ring, single bond, and double bond). In case of atom and bond composition vector will have dimension of nine.

##### Residue property

This module generates binary profiles for each residue in the amino acid sequence, which apprehends the particular physicochemical properties for each input sequence. It allows the user to choose the desired physicochemical property. It has provision to select any of 22 of standard physicochemical properties.

##### AAIndex

This sub-module allows the user to compute the binary profiles corresponding to the desired AAindices, for amino acid sequences. It enables the user to enter the desired indices out of defined 566 amino acid indices, which can be referred from https://www.genome.jp/aaindex/AAindex/list_of_indices. If user select 10 AAindices then each amino acid will be presented by a vector of dimension 10. In case protein have 100 residues then it will be represented by a vector of dimension 1000 (100 × 10).

#### 3.1.3. Evolutionary Information

This module captures the evolutionary information, which plays a significant role in defining the functionality of the peptide or protein. This provides more information than single sequence. Thus, it is an essential representative of a peptide or protein. This module is further sub-divided into four sub-classes:

##### Generation of PSSM

Position-specific scoring matrix can be generated for single sequence provided by user. It uses the PSIBLAST software to generate PSSM profiles. It captures the spatial information for evolutionary conservation of amino acids. It gives the output in the form of PSSM matrix.

##### Normalization of PSSM

In this module user has to provide the PSSM matrix as input for normalization. Even he can use the above mentioned method to generate PSSM matrix or can simply provide matrix generated by somewhere else. Further user can opt any of the four different methods for normalization such as pssm_n1, pssm_n2, pssm_n3 and pssm_n4.

##### Composition of PSSM

In this sub-module user can calculates the composition of PSSM corresponding to each input PSSM matrix. This method gives output in the form of 20X20 matrix.

##### Profile of PSSM

Using this module user become enable to calculate the normalized position-specific scoring matrix using psi-blast against databases. Here user has to provide fasta file sequence as input. This method gives output in the form of matrix of N × 21 for the peptide or protein sequence of length N. It also provides user to generate profiles for different portions of the peptide or protein sequences.

#### 3.1.4. Structure Descriptors

This module is meant to compute a wide range of structure-based features and encompasses four sub-modules:

##### Fingerprint

This module allows the user to calculate the fingerprint descriptors for the input PDB structure. These fingerprints are calculated using PaDEL software (Yap, 2011). This result in the vector of size 14,532.

##### SMILES

This module provides the Simplified Molecular Input Line Entry System (SMILES) notation for the PDB structure, which can be further use to generate structure-based features. These SMILES are generated using open babel software and provide a line notation for representing peptide or protein structure.

##### Surface Accessibility

This module allows the user to calculates the relative surface accessibility of the input PDB structure, using NACCESS software. The output exhibits the surface accessibility for each residue in the peptide or protein sequences. It ranges from 0-100 percent.

##### Secondary Structure

This module generates the secondary structure of the peptide or protein at the residue level. It calculates the percent of average secondary structure element present in the input PDB file. These features are calculated using DSSP software (Touw *et al.*, 2015).

#### 3.1.5. Pattern Generation

This module is meant to generate a wide range of pattern-based features. It is essential to generate all fixed-size length patterns when one is interested in residue level annotation. This module focuses on different size patterns for different profiles. This module encompasses five sub-modules:

##### Binary Profile

This module computes the binary profiles for the patterns of protein or peptide sequences. The patterns are generated using the desired window size which is odd numbers. The equal number of X’s are added in the starting and ending of the input sequence to produce equal size patterns. Each letter corresponds to the vector size of 21, hence generate the vector size of N*21 for the sequence of length N.

##### PSSM Profile

This module calculates the PSSM profiles of the patterns of protein or peptide sequences. The patterns are generated in the same way as mentioned above.

##### Standard PhysicoChemical Properties

This module allows users to calculate the desired standard physicochemical properties from the generated patterns of the sequences of desired window size.

##### AA Index

This module allows users to calculate the average desired AA index value for each generated patterns of desired window size.

##### Universal

This module generates patterns of any type of string or matrix such as peptide or protein sequences, secondary structure, PSSM profiles. The patterns are generated in sliding manners and defined window size. It results in the generation of patterns of equal length.

### 3.2. Standalone version of Pfeature

In addition to web-based implementation, we have also developed standalone of Pfeature, suitable to run on local machine of users. Following is brief description of packages available for Pfeature.

#### Pfeature Scripts

More than 120 python scripts have been developed for computing wide range of features, visit GitHub website https://github.com/raghavagps/Pfeature/scripts/ for detail. These scripts can be downloaded as a zip file from https://github.com/raghavagps/Pfeature/blob/master/scripts/Pfeature_scripts.zip. User need to unzip this file to create a folder of python scripts. Summary of these scripts with their function is described in Table 1. User can run these scripts by typing following command “python script.py--h”, which will provide help on usage of python script. As shown in Table 1, aac_X is for computing amino acid composition, it means following scripts and type of compositions, i) aac_wp.py for whole protein, ii) aac_nt.py for N-terminal residues, iii) aac_ct.py for C-terminal residues, iv) aac_rt.py for remaining residues and v) aac_st.py for fragment of proteins. Thus, scripts not only provide facility to compute composition of whole protein but also allows, to compute composition of a particular region of a protein. In case of tripeptide composition, we only allow to compute tripeptide composition using script tpc.py not for different part of proteins because number of features are too high. These scripts take protein/peptide sequences in FASTA format and give output file csv format.

**Table 1:**
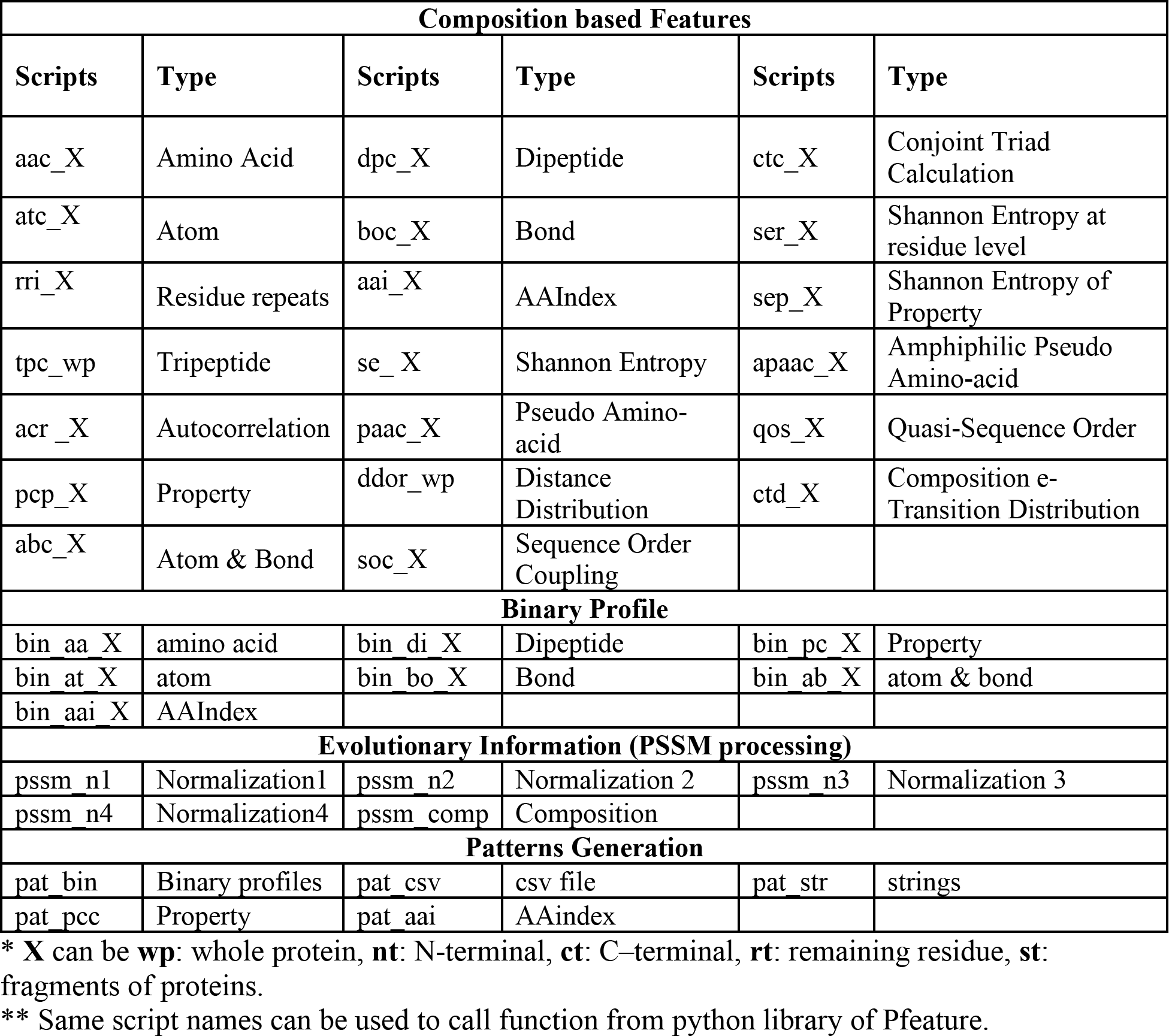
List of python scripts (with extension “.py”) and their corresponding features.

#### Python Libraries

In order to develop method for annotating proteins in python user may wish to call python functions for speed optimization. We have also developed python library of Pfeature, which can be downloaded from https://github.com/raghavagps/Pfeature/blob/master/PyLib/Pfeature.zip. The prerequisite to run the python library is pandas, numpy and python version 3.6 and above. The zipped file can be uncompressed. There is a file inside Pfeature folder named “setup.py”, which helps in installation of library using command “python setup.py install”. This *Pfeature* library consists of 120 functions for descriptor generation. These functions can easily be imported and compute the desired features by passing the required arguments.

#### Pfeature Executables

One of the challenges in computing features of protein is that there are huge number of features. In case, we wish to compute all type of features we need to run more than 100 python script, which is time consuming. We are inspired with the field of chemoinformatics where software compute wide range of chemical descriptors in a single command, for example PaDEL compute more than 10000 chemical descriptors in a single command. Thus, we have developed a standalone version of Pfeature which can compute thousands of protein descriptors in a single command. To cover maximum user, we have developed executables for Windows, Ubuntu, Fedora, MacOS and Centos based operating system. These executables can be downloaded from https://github.com/raghavagps/Pfeature/tree/master/exec/.

### 3.4 Comparison with other resources

Comparison of newly developed software with existing software give insight about its significance (Table 2). Only Pfeature is available in all three format of webserver, standalone, executables and library. Pfeature computes nearly 144 type of protein features where other methods allow 14 to 53 type of features. Thus features computed by Pfeature is nearly three times of features computed by any existing software. Pfeature includes all the types of protein features computed by existing methods and have more than 90 types of additional features. In order to compare different types of features computed by different methods including Pfeature, we show different types of features and supported software in Table 3. Following are unique descriptors not supported in any existing software package.

**Table 2:** Comparison of Pfeature with other software in term of total features and availability.

| Software | Types of protein descriptors | Webserver | Standalone/Scripts | Library |
| --- | --- | --- | --- | --- |
| Pfeature | 144 | Yes | Yes | Yes |
| iFeature | 53 | Yes | Yes | Yes |
| PyBioMed (PyProtein) | 14 | No | Yes | Yes |
| PyDPI | 52 | No | Yes | Yes |
| PROFEAT | 51 | Yes | No | No |

**Table 3:**
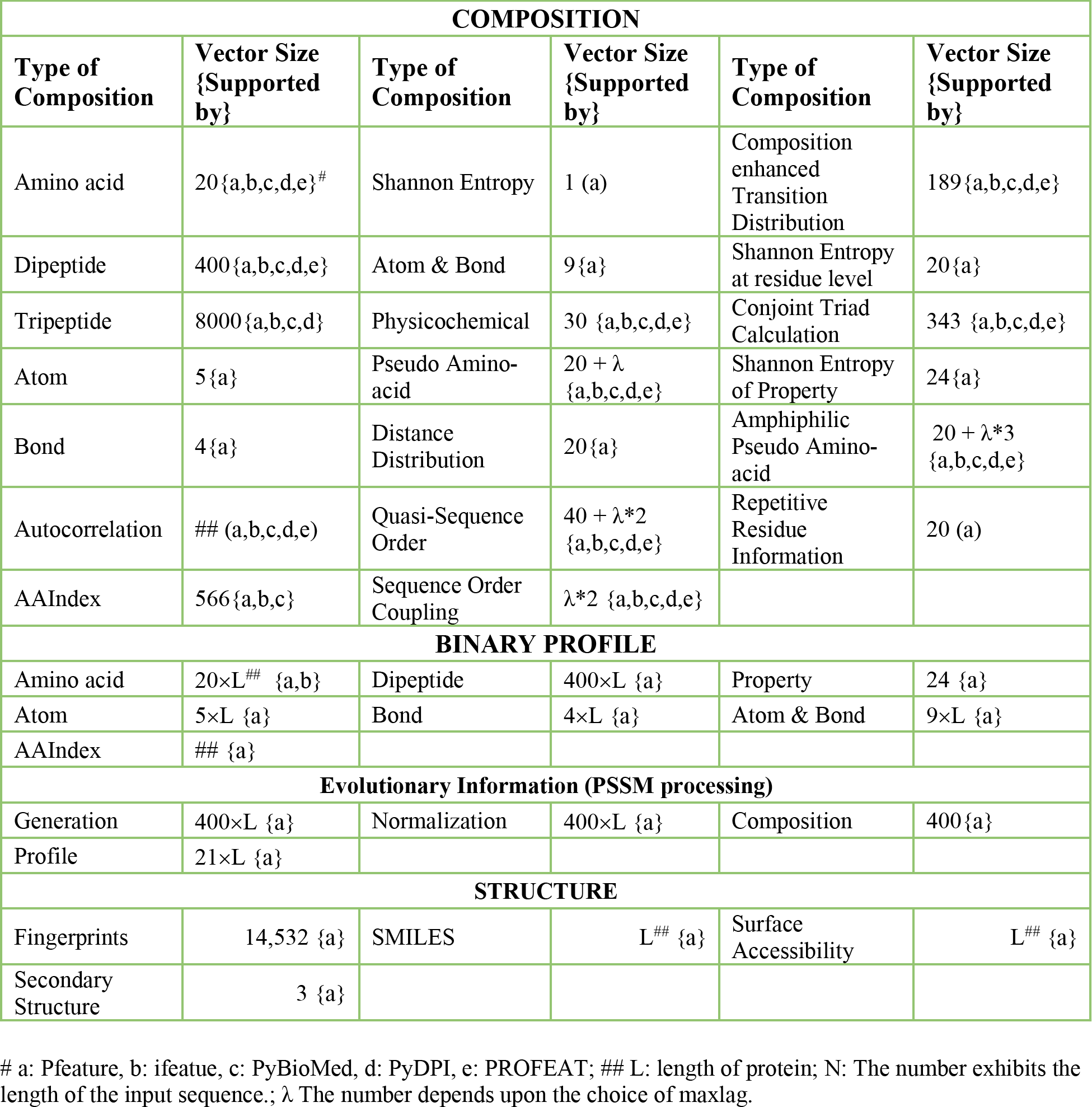
Comprehensive Comparison of Pfeature with existing software/platform in term of different type of features.

As shown in Table 3, following composition based modules has been introduced in this study; atom, bond, atom+bond, Shannon entropy, residue repeats and distance distribution. Repetitive Residue Information computes the repetitive residue information for a peptide sequence. In more simpler words, it actually measures the continuous presence of amino acids in a protein or peptide sequence. Distance distribution function calculates the distribution of residues and considers the distance from the terminus residues of the peptide or protein sequence. In addition, almost all binary profile-based features are only supported by Pfeature. Pfeature incorporated the function which is used to compute amino acid binary profiles, dipeptide binary profiles, a binary profile corresponding to atomic and bond composition of each amino acid residue of the peptide sequence. Pfeature also support PSSM profile-based features that is required to capture evolutionary information, which are not supported by existing tools.

## 4. Discussion

In past, several web-servers and standalone packages have been developed to extract features from protein and peptide sequences. PROFEAT is one of the first web servers developed in 2006, for computing structural and physicochemical features of proteins and peptides from amino acid sequence (Li *et al.*, 2006). Its updated versions were published in 2011 and 2016 that incorporate new network descriptors for protein-protein and protein-small molecule interactions, segment descriptors for local properties of protein sequences, topological descriptors for peptide sequences and small molecule structures (Rao *et al.*, 2011; Zhang *et al.*, 2017). Though, our Pfeature have more protein descriptors than PROFEAT but we do not have many network descriptors integrated in PROFEAT. Thus, Pfeature is complementary to PROFEAT but not alternate or replacement. Similarly, PyBioMed is a recently developed python library with the capability of computing features of biological molecules and interaction samples (Dong *et al.*, 2018). A module named “PyProtein” is developed in PyBioMed which is used to calculate protein features (structural and physicochemical properties). PyProtein module calculates 14 features from five groups. Pfeature also integrate all these descriptors and many more protein descriptors but it does not have modules for computing DNA and chemical descriptors, integrated in PyBioMed. PyDPI is feature rich python based standalone that computes 52 types of protein features from six feature groups (Cao, Liang, *et al.*, 2013). Recently, one of the powerful package iFeature has been developed that compute a comprehensive spectrum of 18 major sequence encoding schemes that encompass 53 different types of feature descriptors. It also integrates 12 different types of commonly used feature clustering, selection, and dimensionality reduction algorithms. These features are important for developing machine learning techniques-based models for predicting function of biomolecules.

In summary, number of software package has been developed to compute descriptors of biological & chemical molecules. Each package has unique set of features that complement other existing packages. Aim of developing Pfeature is to complement the existing software so user may get more options in the field of protein bioinformatics. Our major focus is on protein annotation that includes chemically modified protein and peptides. We are particularly interested in developing features for computing therapeutic properties of peptide and annotation of protein at residue level. Despite tremendous advancement in the field of protein bioinformatics, a wide range of protein descriptors used in past has not been integrated in existing software packages. Residue level annotation of a protein is highly desirable to understand function of a protein at residue level. In addition, these softwares do not have facility to compute composition of a specific-region of protein. In order to facilitate users, we have developed Pfeature that integrate most of descriptors described in literature as well as few novel descriptors. In summary, it is a comprehensive, easy-to-use python package which computes a large number of features, and allows users to calculate various features of protein/peptide based on the sequence, structure and physiochemical properties. As far as we know, Pfeature is the first python package that calculates the several descriptors and gives the information of not only at functional level but also at the residue level. It is the first toolkit for combined feature calculation at the functional level, residue level and also from the various profiles such as binary profiles and PSSM profiles. We believe that freely available web service “https://webs.iiitd.edu.in/raghava/pfeature/” will be very useful to the biologist, with a limitation of programming knowledge. In addition, free-availability of python library, executables as well as source code will help the researcher to compute a wide range of protein and peptide feature from their sequence and structure on a larger scale.

## Acknowledgements

Authors are thankful to funding agencies Department of Biotechnology (DBT) and Department of Science and Technology (DST-INSPIRE) and Council of Scientific and Industrial Research (CSIR), Govt. of India for financial support and fellowships.

## Author contribution

SP, AP, AL, CA, DK, AD and GM wrote all the scripts. HK, SP, AD, AL, CA, DK, SSU, PA, RK and VK developed the web interface. NS, SJ, AL and CA prepared the manual. SSU, SP, AP, NS and AD prepared the first draft of manuscript. SSU and GPSR prepared the final version of manuscript. GPSR conceived the idea and coordinated the entire project.

